# Fotomics: Fourier transform-based omics imagification for deep learning-based cell-identity mapping using single-cell omics profiles

**DOI:** 10.1101/2022.07.08.499309

**Authors:** Seid Miad Zandavi, Derong Liu, Vera Chung, Ali Anaissi, Fatemeh Vafaee

## Abstract

Different omics profiles, depending on the underlying technology, encompass measurements of several hundred to several thousand molecules in a biological sample or a cell. This study develops upon the concept of “omics imagification” as a process of transforming a vector representing these numerical measurements into an image with a one-to-one relationship with the corresponding sample. The proposed imagification process transforms a high-dimensional vector of molecular measurements into a two-dimensional RGB image to enable holistic molecular representation of a biological sample and to improve the classification of different biological phenotypes using automated image recognition methods in computer vision. A transformed image represents 2D coordinates of molecules in a neighbour-embedded space representing molecular abundance and gene intensity. The proposed method was applied to a single-cell RNA sequencing (scRNA-seq) data to “imagify” gene expression profiles of individual cells. Our results show that a simple convolutional neural network trained on single-cell transcriptomics images accurately classifies diverse cell types outperforming the best-performing scRNA-seq classifiers such as support vector machine and random forest.

## Introduction

Modern data-driven biology relies on the interpretation of large-scale molecular measurements (known as *omics* data) to understand and predict biological *phenotypes*, such as the characteristics of an organism or the state of an individual cell. Next-generation sequencing (NGS) biotechnologies can now comprehensively measure different types of molecules (e.g., DNA, RNA, and proteins) at a high-speed and low-cost, leading to an unprecedented generation of *omics* data from different biological samples or across thousands of individual cells. The ‘omics’ notion indicates that nearly all instances of the targeted molecules are measured in the assay providing holistic views of the biological system [1]. For instance, *transcriptomics* data derived from RNA-sequencing (RNA-seq) technologies measure the quantity of over 20,000 RNAs in a biological sample at a given moment. Assay for Transposase-Accessible Chromatin using sequencing (ATAC-seq) measures chromatic accessibility representing the level of physical compaction of chromatin across the genome.

Conventional RNA-seq experiments measure the *average* RNA expressions from large populations of cells within a sample or tissue section. Such average measurements, however, may obscure critical differences between individual cells within these populations [2]. In contrast, *single-cell* RNA sequencing (scRNA-seq) provides transcriptomics profiles of thousands of individual cells providing unprecedented opportunities to characterise the cellular composition of complex tissues. Analysis of single-cell sequencing data has taken the front seat in deriving useful insights into cellular functions and identification of cell sub-types with diverse applications across biological disciplines, including developmental biology, neurology, oncology, immunology, and Infectious disease.

A common step in analysing single-cell data involves the identification of cell populations present in a given sample. This task is accomplished either via unsupervised clustering [3] (i.e., grouping cells based on similarities in their expression profiles followed by cell annotation, which involves manual inspection of cluster-specific marker genes) or via supervised classification enabling automated cell identification and annotation [4]. The latter circumvents the cumbersome and time-consuming step of manual cell annotation and has increasingly become popular due to the growing number of annotated scRNA-seq datasets available for training classifiers [5]. A recent study has benchmarked over 20 classification methods (including both single-cell-specific and general-purpose classifiers) to automatically assign cell identities across several scRNA-seq datasets of different sizes, technologies, and complexity and demonstrated that the general-purpose support vector machine (SVM) classifier is the best performer, overall [4].

However, SVM, similar to other traditional classifiers, requires a feature selection step prior to training to identify informative features (e.g., differentially-expressed genes) or latent variables (e.g., features extracted via dimensionality reduction) as predictors of the classifier. The choice of feature selection often significantly affects classification performance and further complicates the application of a classifier for cell identification [6]. Additionally, while SVM is proven to be a powerful scRNA-seq classifier, it shows varied performance across different datasets and fails to correctly determine cells in complex populations which incorporate several closely-related cellular subtypes [7]. This signifies the intricacy of expressional patterns underlying different cell types and the need for an end-to-end learning platform that can decipher subtle differences among cell types.

Deep artificial neural networks are widely acknowledged for their ability to perform *automatic feature extraction* from raw data. Hence, a deep learning-based model has the capacity to develop a single-step phenotype prediction procedure where omics data would be directly used to classify the outcome without prior feature selection. The absolute superiority of convolutional neural networks (CNNs) in image classification has been widely acknowledged but not yet sufficiently leveraged for classifying cell types based on single-cell omics data. Intuitively, CNN’s convolution operators extract local spatial features from an input image. The local information is then combined, via pooling aggregations, to higher-order special information from a large region of an image which would be eventually used to distinguish among different image types. Therefore, the utility of CNN is in the automated extraction of local and global special features from input images where adjacent pixels are interrelated and share similar information.

Converting an omics numerical vector to an image requires the placement of spatially coherent pixels in local regions to incorporate interrelationships and local/global patterns in molecular profiles into phenotype prediction. In other words, an arbitrary arrangement of pixels or features in a 2D coordinate negatively impacts the performance of feature extraction and classification, defeating the purpose of imagification. In this study, we presented a method to convert a vector representing single-cell sequencing measurements to an image (matrix data keeping spatial image information), referred to as ‘omics imagification’ combined with CNN-based classification for cell-type identification.

Here, we proposed a new method inspired by discrete Fourier transformation [8] in signal processing referred to as *Fotomics* (*<u>Fo</u>urier transformation-based o<u>mics</u>*) imagification for cell identity mapping using single-cell sequencing data. Fotomics is developed based on the idea that a cell is a biological system whose characteristics can be understood by a collective quantification of pools of molecules (i.e., omics). In a nutshell, upon a Fourier transformation of cellular omics profile, Fotomics maps each feature into a complex Cartesian plane wherein proximities of features reflect their similarities and, thus, produces images with special relationships among pixels. We demonstrated that Fotomics enhances prediction performance compared with the state-of-the-art image conversion method and best-performing classifiers for automated cell-type identification using scRNA-seq data.

## Related Works

While converting omics data to images is a relatively new concept in the field of bioinformatics, former studies have used different CNN architectures to classify biological or clinical samples based on omics measurements – often gene expression data – converted into two-dimensional images.

Kovalerchuk et al. [9] proposed a generic algorithm, referred to as CPC-R, to convert non-image data to images through pair values mapping. In a nutshell, the proposed algorithm splits an *n*-dimensional feature vector *X* = (*x*_1,_ *x*_2, …,_*x*_*n*_) to consecutive pairs (*x*_1,_ *x*_2_), (*x*_3,_ *x*_4_), …(*x*_*n*−1,_ *x*_*n*_). It then sets up a cell size to locate pairs in a 2D image where each cell can consist of a pixel or dozens of pixels. Finally, each pair (*x*_*i*_, *x*_*i*+1_) is located at the corresponding cell coordinate pair of the image. This algorithm, however, assumes an initial fixed and meaningful ordering of features in an *n*-dimensional vector which is not a valid assumption in omics data.

Further, Lyu and Haque [10] converted RNA-Seq data into 2-D images and used a CNN to classify 33 tumour types obtained from The Cancer Genome Atlas, TCGA (www.cancer.gov/tcga). The 2D images were constructed by first ordering genes based on the chromosome number and then reshaping the corresponding 10381×1 array into a 102 × 102 matrix by adding zeros to the last line of the matrix. Finally, the expression values were normalised to [0, 255], representing a grayscale image. Here, the ordering of genes is based on the assumption the adjacent genes on the chromosomes are more likely to interact with each other. However, the currently accepted models of transcriptional regulation support long-range interactions between proximal as well as distal regulatory regions on the genome (e.g., promoter–enhancer interactions) [11], weakening the assumption underlying the proposed imagification procedure.

Later, Lopez et al. [12] proposed a methodology to transform gene-expression vectors into images wherein the relative position of the genes is driven by their molecular function. The KEGG ontology database [13] has been used to query the BRITE hierarchies comprising the genes of the input vector. A treemapping [14] algorithm has been used to display the hierarchical ontology data onto images by using nested rectangles, representing KEGG functional categories and sub-categories that eventually contain genes. The proposed approach was applied to RNA-seq data obtained from TCGA dataset, and the corresponding images were trained using a CNN combined with transfer learning to predict patient survival.

Additionally, Sharma and Kumar [15] proposed three methods to transform a 1-D vector to a 2-D image with associations among the fields/pixels to be processed by CNN. The methods were used to predict breast cancer using numerical vectors of patients’ clinical information obtained from Wisconsin Original and Diagnostic Breast Cancer datasets from UCI library [16] The first method involves using the bar graph to visualise every feature of the dataset. The height is normalised to create a square image. The order of the bars is crucial for CNN to depict patterns and was determined based on the similarity of features. This was performed by first constructing a covariance matrix and converting values to ranks, and then determining an optimal order of bars based on their ranks using dynamic programming or a metaheuristic optimisation algorithm [17]. The second method uses the normalised distance matrix of size *d* × *d*, where *d* is the number of features and matrix elements are differences between the corresponding features normalised between [0 – 1]. The third method combines the other two strategies by creating a three layers image of size [3*d* × 3*d*] where a copy of numerical data, the bar graphs and a normalised distance matrix stack on top of each other. The third method was shown to outperform the two other methods. These methods were not applied only applied to convert clinical feature vectors of size ≤32 variables and are not feasibly applicable to large-scale omics data comprising several thousands of variables.

Among these methods, DeepInsight, introduced by Sharma et al. [18], is the most acknowledged algorithm for transforming non-image data into images applied to different data types, including RNA-seq, vowels, text, and synthetic data for the purpose of classification using CNNs. DeepInsight uses a dimensionality reduction technique such as t-SNE [19] across samples to obtain a 2D plane where the points in this Cartesian plane define the locations of the features (e.g., genes in RNA-seq data), followed by the convex hull algorithm to find the smallest rectangle containing all the points, and rotation to frame a horizontal or vertical image to input into a CNN architecture. Finally, the Cartesian coordinates were converted to pixels, and for each sample, the feature values (e.g., gene expressions) were mapped to the pixel locations. Later, the DeepInsight authors developed DeepFeature [20], which relies on DeepInsight-obtained images to select important features from non-image data using CNNs, further corroborating the utility and subsequent applications of imagification.

In this paper, we propose a novel image transformation method to convert non-image omics data into images using Fast Fourier Transform (FFT) to map features onto a complex Cartesian plane. The proposed method was applied to single-cell RNA-seq data for cell-type classification using CNN models. The performance of the proposed method was compared with DeepInsight, as well as traditional classifiers used for cell identity mapping using scRNA-sequencing data.

## Materials and Methods

### Overview of Fotomics

**Figure 1** illustrates the overall workflow of Fotomics imagification. Assume an input *n*×*m* matrix representing molecular measurements of *n* features (e.g., genes) across *m* cells or biological samples. Here, we consider each column as an *‘omics signal’* of length *n*, comprising a series of molecular measurements (ordered in any random arrangement) from a cell or sample.

**Figure 1.**
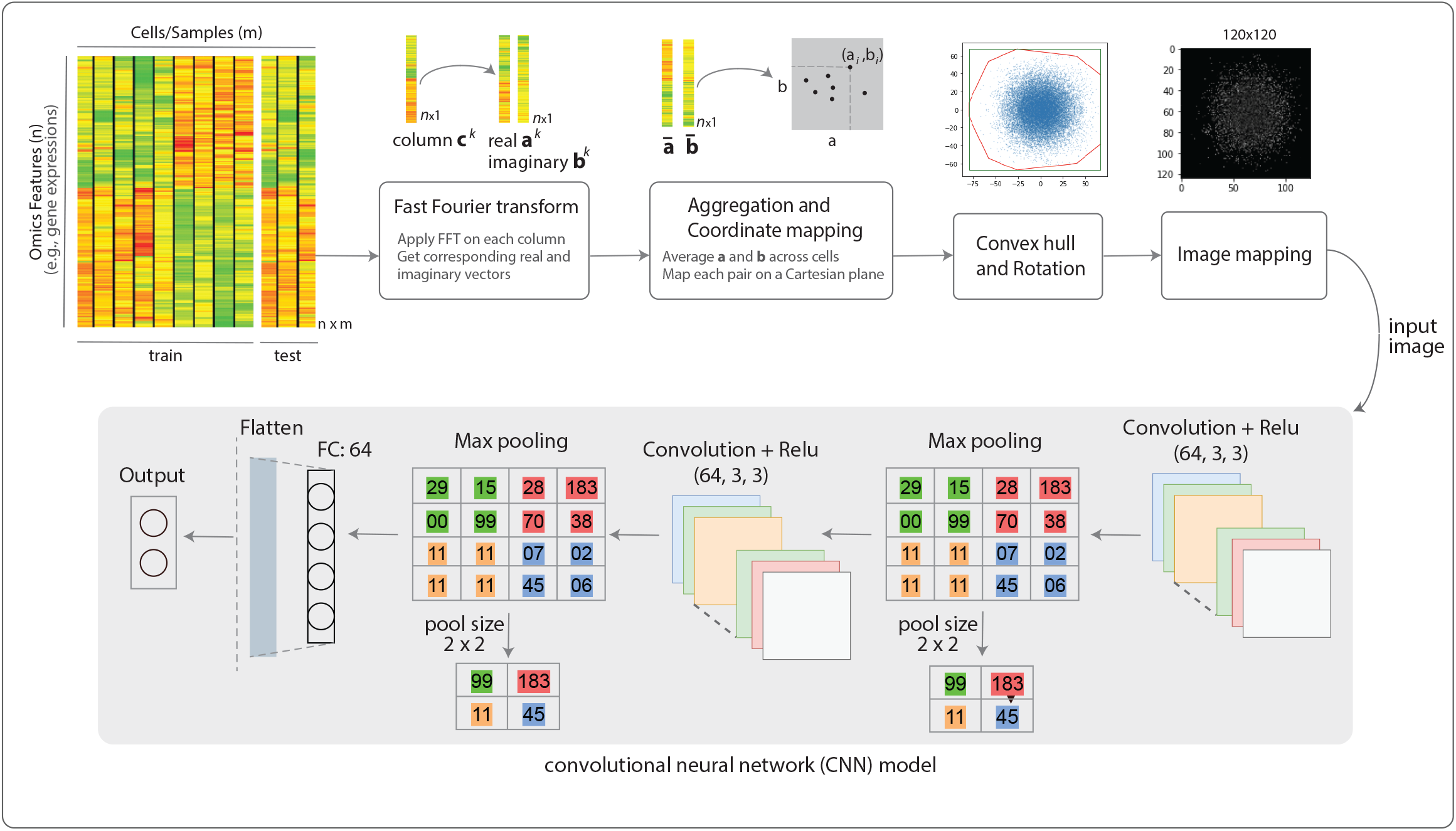
Fotomics imagification workflow. The input omics profile (log-transformed) is split into test and train. The train set undergoes the illustrated procedure (also detailed in Figure 2 pseudocode). The image coordinates obtained from the training set will be also used to convert test samples into images.

After a preprocessing step (e.g., normalisation to account for unwanted variations across samples), each column undergoes the discrete Fourier transformation (DFT), a mathematical approach for transforming a finite data sequence into the frequency domain to reveal hidden patterns and periodicities. The fast Fourier transform (FFT) [21] is a highly efficient procedure to compute the DFT of a data sequence. FFT of a sequence of length *n* returns an equally sized vector of complex numbers, each represented as

*a* + *bi*, where *a* and *b* are the real and imaginary parts, representing the magnitude and phase of the signal, respectively. A complex number can be represented in a Cartesian plane (*a-*axis versus *b*-axis), allowing a geometric interpretation of the complex number.

Accordingly, upon applying FFT to each column *k*, we obtain two vectors of size *n* representing real values 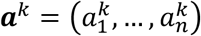 and imaginary values 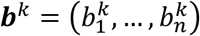. These vectors are averaged across samples to get ***ā*** = (*a*_1,_ …, *a*_*n*_) and 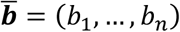, where 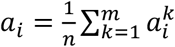 a similar equation holds for *b*_*i*_. We then map ***ā*** and 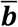 on a Cartesian plane where point *i* with coordinates ⟨*a*_*i*_, *b*_*i*_⟩ corresponds to feature *i*. The proximity of points within this Cartesian plan unveils the underlying associations or similarities between different features across cells. The output of this process is a *reference* map where the location (i.e., Cartesian coordinates) of each feature is determined. As previously suggested [18], [20], we then applied a convex hull algorithm to find the smallest convex set that contains the points within the Cartesian plain, followed by rotation to frame the plane vertically or horizontally to comply with the CNN input orientation. Finally, we considered a 2D image with predetermined pixel dimensions and mapped each point within the coordinate into the respective pixel location. The pixel intensity is determined by the mean expression values of the features in the same location. Pixel dimensions need to be carefully determined to avoid obscuring useful information while considering computational resources. When the picture is very small, multiple distinct features get mapped into an identical pixel location, and the visual representation could be inaccurate when compared to the number of features available. **Figure 2** Shows the pseudocode of the Fotomics algorithm.

**Figure 2.**
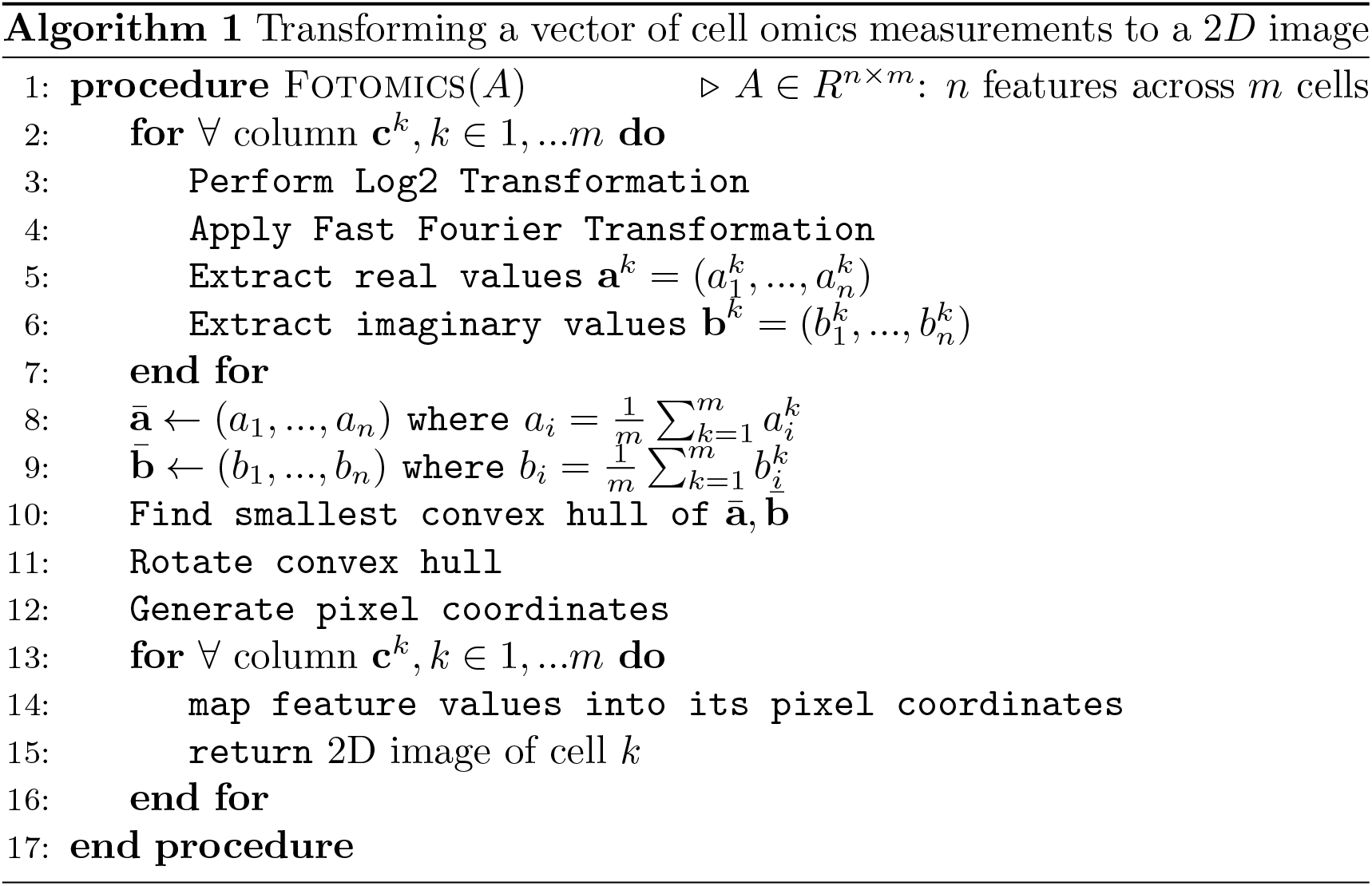
Fotomics pseudocode.

While Fourier transformation assumes an ordering of input features, it has been proven by Lanczos and Gellaithe [22] that Fourier transformation can be used to search for hidden periodicities in ‘random sequences’. We also showed previously [7] that Fourier-based extracted patterns are independent of the initial random ordering of the omics features. Therefore, any initial random ordering of the features should be able to map relevant features in close proximity on the complex Cartesian plane; the ordering should remain the same across cells or samples.

### CNN architecture

The convolutional neural network (CNN) architecture illustrated in Figure 1 includes a combination of a convolution layer, plus a ReLU activation function, then a max-pooling layer with a pool size of 2×2. There are two of these combination sequences, followed by a fully connected and a flattened layer. To avoid overfitting, we have used multiple strategies, including train-validation-test splitting, simplifying the architecture, and reducing the depth of CNN, early stopping, learning rate reduction on the plateau, L1/L2 regularisation, and dropout regularisation. The reported accuracies are on the test set and have been consistent with the training loss demonstrating that overfitting was avoided to a sufficient degree.

### Data collection

A total of 14 datasets were obtained from a curated publicly available scRNA-seq dataset maintained by the Hemberg Lab [23]. Each dataset includes labels for predicted cell types, which were used as the ground truth for classifier training and performance evaluation. These datasets are selected to include a wide range of protocols, tissue types, organisms, and dataset sizes, as detailed in Table 1. Given the increasing understanding that scRNA-seq data generated with protocols that employ Unique Molecular Identifiers (UMIs) have different properties from read-based data [24], we included a balanced combination of UMI-based datasets (n=7) and read-based datasets (n=7).

**Table 1.** Summary of scRNA-seq datasets used to evaluate Fotomics. Databases are alphabetically sorted by their name; the associated numbers were used to reference them in relevant figure(s).

| No | Dataset | Protocol | Accession | Species | Tissue type | Cells | Genes | Cell Types | UMI/Read |
| --- | --- | --- | --- | --- | --- | --- | --- | --- | --- |
| 1 | darmanis [ref] | SMARTer | GSE67835 | Human | Brain | 466 | 22088 | 9 | Read |
| 2 | deng reads [36] | Smart-Seq2 | GSE45719 | mouse | embryo | 268 | 22431 | 6 | Read |
| 3 | Fan [37] | SUPeR-seq | GSE53386 | mouse | embryo | 66 | 26357 | 6 | Read |
| 4 | heart 10X [38] | Chromium | GSE132042 | mouse | heart | 654 | 23433 | 7 | UMI |
| 5 | liver 10X [38] | Chromium | GSE132042 | mouse | liver | 1924 | 23433 | 3 | UMI |
| 6 | manno mouse [39] | STRT-Seq | GSE76381 | mouse | brain | 2150 | 24378 | 32 | UMI |
| 7 | marrow 10X [38] | Chromium | GSE132042 | mouse | marrow | 4112 | 23433 | 9 | UMI |
| 8 | pancreas FACS [38] | Smart-Seq2 | GSE132042 | mouse | pancreas | 1961 | 23433 | 10 | Read |
| 9 | tasic reads [40] | SMARTer | GSE71585 | mouse | brain | 1679 | 24150 | 18 | Read |
| 10 | Trachea FACS [38] | Smart-Seq2 | GSE132042 | mouse | trachea | 1391 | 23433 | 5 | Read |
| 11 | yan [41] | Tang | GSE36552 | human | embryo | 90 | 20214 | 6 | Read |
| 12 | zeisel [42] | STRT-Seq | GSE60361 | Mouse | brain | 3005 | 19972 | 9 | UMI |
| 13 | zhengmix4eq [43] | GemCode | SRP073767 | human | immune | 3994 | 15568 | 4 | UMI |
| 14 | zhengmix4uneq [43] | GemCode | SRP073767 | human | immune | 6498 | 16443 | 4 | UMI |

### Data preprocessing

For each dataset, features with zero values across all cells were filtered out. Following a 5-fold cross-validation procedure, 20% of cells or samples were randomly picked and held out as the test set, and the remaining 80% of cells were used for training (and hyperparameter optimisation), and the process was repeated 5 times on disjoint test sets. For cells in the training set, read count values were normalised to account for unwanted and technical variations in measurements across samples. We implemented two widely used normalisation methods, namely count per million (CPM) [25], and minimum-maximum scaling on CPM data (MinMax) [26].

The input data (i.e., the training data) was processed by log transformation and scaling as formulated in **Equations 1 – 3**, where *X*(*i*, :) represent row *i* comprising the values of feature *i* across samples in the training data, and 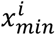 is the minimum value of *i*-th feature across samples. *X*′ and *X*′′ are log transformed and scaled form of data, respectively. The processed parameters obtained from the training set were used to normalise the test set to avoid any potential leakage of information from the test to train set (consider a scenario when the entire dataset is normalised prior to test-train splitting wherein the training samples are normalised based on the information contained in test samples causing test-to-train information leakage).

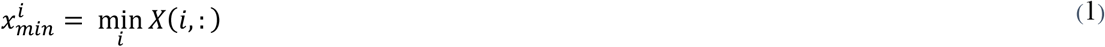

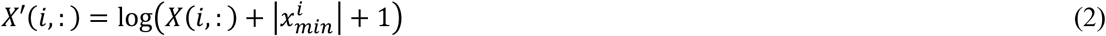

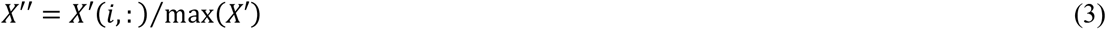

### Image production

The processed data is then undergone fast Fourier transformation to identify the 2D coordinates of each feature as previously described. Then, the convex hull is carried out to find the minimum convex that encompasses all the features on the coordinate plane followed by rotation as formulated in **Equation 4**, where [*a*_*i*_ *b*_*i*_]^*T*^ is the resulting images after FFT transformation, from which *a*_*i*_ and *b*_*i*_ are real and imaginary numbers, respectively. [*a*_*ri*_ *b*_*ri*_]^*T*^ is the rotated image planes.

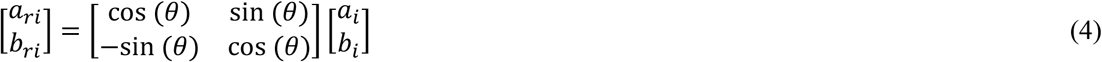

**Equations 5 – 6** formulate the coordination of each feature based on the pre-determined pixel size in the rotated plane, where *x*_*c*_ and *y*_*c*_ are computed pixel coordinated, *P*_*x*_ and *P*_*y*_ are pre-determined pixel size. Pixel size should be large enough while considering the hardware capability and availability of computational resources.

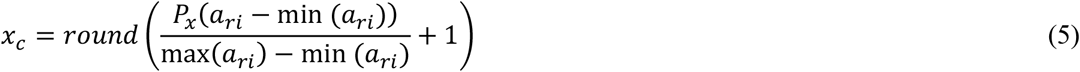

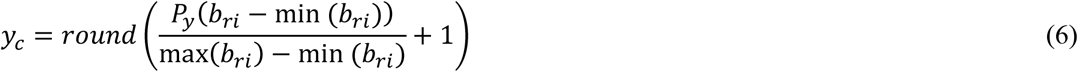

## Results and Discussion

The Fotomics imagification pipeline, as depicted in **Figure 1** and demonstrated in **Figure 2** pseudocode, was applied to 14 scRNA-seq datasets (Table 1) with varying characteristics in terms of the sequencing protocols, tissue, organism, and *complexity* level as defined previously [27]. **Figure 3A** shows examples of images from 4 different datasets, each representing an arbitrarily chosen cell within a dataset. **Figure 3B** represents the per-pixel density of images across the entire dataset (i.e., scaled mean of h pixel value across all samples).

**Figure 3.**
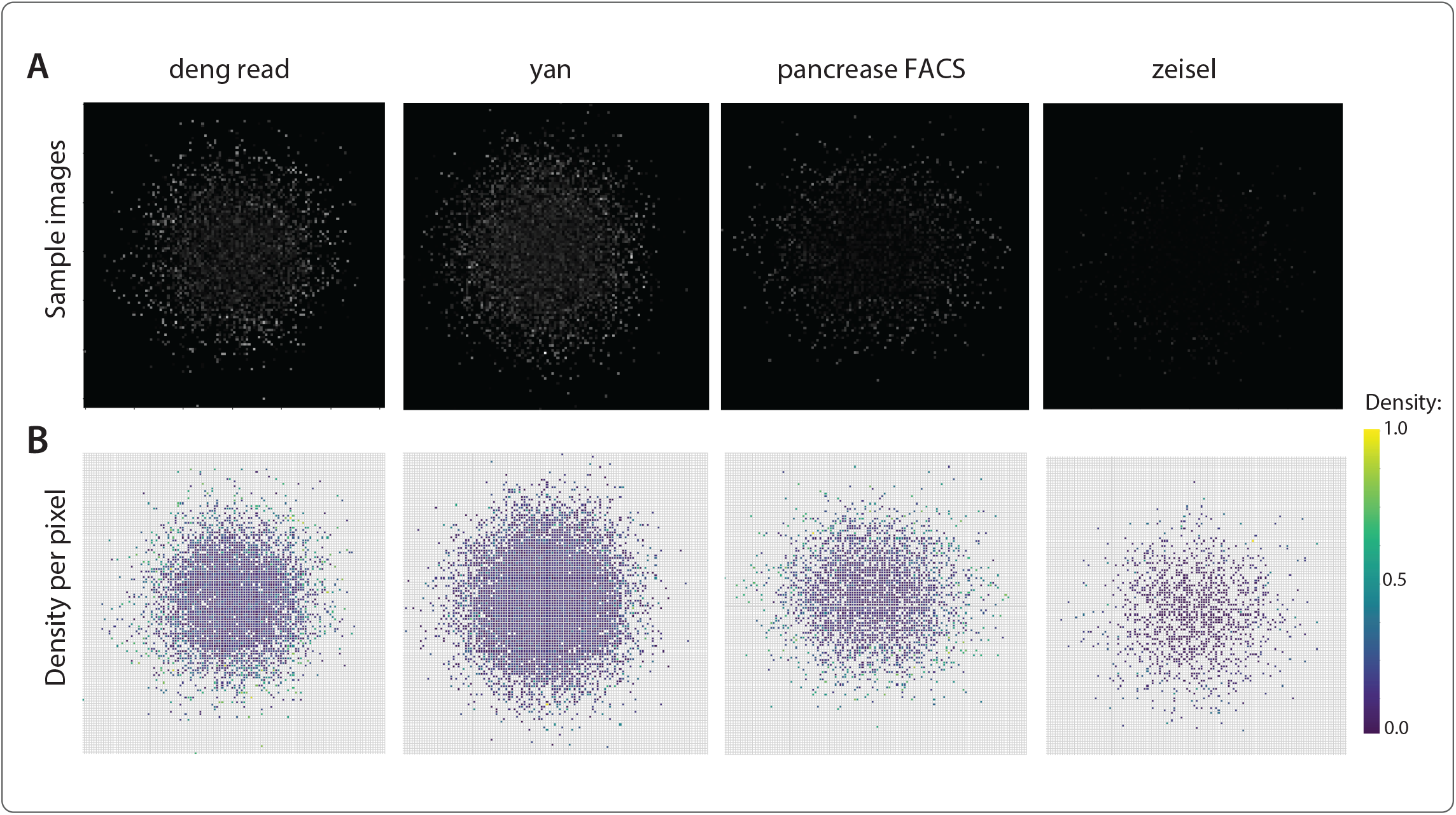
A) Examples of images from four different datasets, each representing an arbitrarily chosen cell within a dataset, and B) per-pixel density of images across the entire dataset.

The utility of Fotonics image transformation for cell type prediction was compared with conventional classification based on numeric features as well as a CNN-based classification based on a former image transformation technique. Recently, Abdelaal et al. [4] have benchmarked 22 classification methods that automatically assign cell identities, including single-cell-specific classifiers and general-purpose classifiers, on a large number of publicly available single-cell RNA sequencing datasets. Upon this comprehensive benchmarking, they have shown that a general-purpose support vector machine (SVM) classifier [28] has overall the best performance across the different experiments. We, therefore, considered SVM as the state-of-the-art classifier for cell identity mapping using scRNA-seq data to compare our model with. We also reported the prediction performance using K-nearest neighbours (KNN) [29] and random forest (RF) [30] classifiers. Additionally, we compared Fotomics with DeepInsight [20], which is the best-performing method amongst formerly developed non-image-to-image transformation approaches, as per a comprehensive benchmarking study under progress by our team. The image pixel dimensions were set to 124×124. For a fair comparison, a similar CNN model was used to train upon Fotomics and DeepInsight-generated images.

Each dataset was trained after a train-test split using a 5-fold cross-validation procedure. Samples were normalised using *cpm* and *min-max* methods, as previously explained. We also used raw, unnormalised data to assess the effect of preprocessing on the downstream performance. The accuracy and macro-averaged F1-score (i.e., unweighted mean of all the per-class F1 scores) were reported upon predicting the cell types of test samples.

Overall, the FFT approach has demonstrated a competitive and consistent result across all 14 datasets with different data preprocessing methods. Three traditional machine learning models (KNN, RF, and SVM) perform poorly compared to CNN models on omics images, both in terms of a higher prediction performance on the test set as well as smaller standard deviations representing the performance robustness, as shown by accuracy and F1-score box plots in **Figure 4**. For example, the RF model did not perform well in the ‘zeisel’ dataset with an F1 score of 22% on raw data, which was significantly lower than others, FFT (89%), KNN (85%), and SVM (68%). Similarly, the SVM model did poorly in the ‘liver-10X’ dataset with an F1-score of 50%, whereas the rest achieved F1-scores of >92%. The KNN model in the ‘darmanis’ dataset had the lowest F1 score of 17%. On the other hand, the FFT model has shown a high level of consistency and outperformed its peers across multiple datasets.

**Figure 4.**
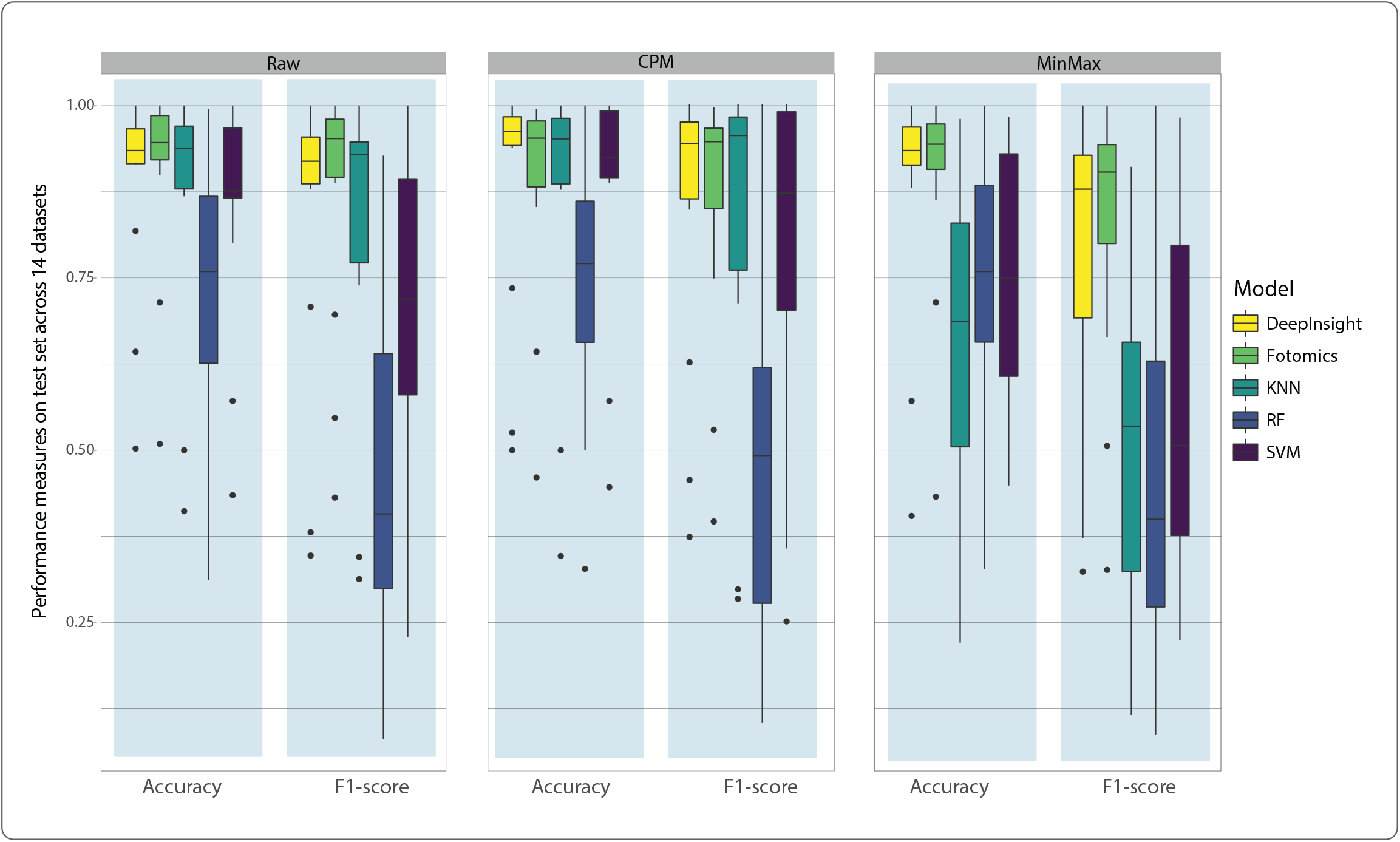
Boxplots representing the distribution of classification accuracy and F1-score on the test set (20% randomly held out cells) comparing conventional classifiers (SVM, RF, and KNN) on vectors of gene expression values and CNN on transcriptomics images of cells generated by Fotomics and DeepInsight. On top of the heatmaps, t-test p-values represent the significance of prediction enhancement by Fotomics compared to other competing techniques.

Fotomics average performance was slightly higher than the DeepInsight algorithm. Across the 14 datasets, DeepInsight had average F1-scores of 88% (raw), 78% (cpm), and 77% (min-max), while FFT achieved 90% (raw), 80% (cpm), and 76% (min-max). Nonetheless, the enhancement was not statistically significant (*p*-value of t-test > 0.5, **Figure 5**). However, Fotomics showed clearly higher classification performance on ‘hard’ datasets where either the number of samples is limited (‘fan’ dataset # 3) or the number of cell types is high (‘manno-mouse’, dataset #6), as shown in **Figure 5**.

**Figure 5.**
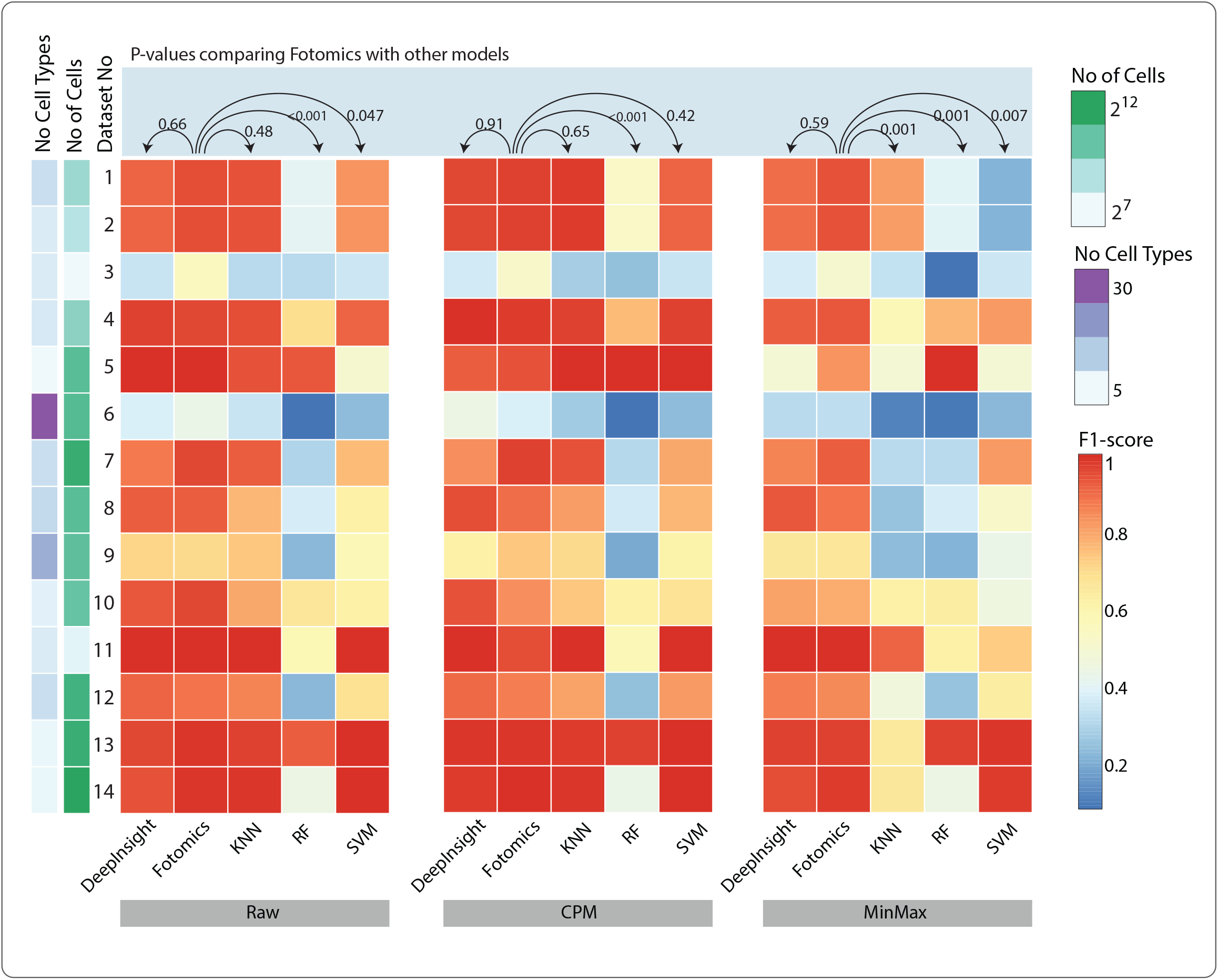
Heatmaps representing the F1-scores of cell-type predictions – for each of 14 scRNA-seq datasets, either as raw counts or normalised with CPM or MinMax – on 20% randomly held out data (test set) after training the CNN model on the remaining 80% of data (training set). Datasets are colour-coded by the number of cell types and number of cells to demonstrate the performance with respect to the complexity of the classification task.

While Fotomics and DeepInsight perform on par across most datasets with respect to classification accuracy, Fotomics is computationally more efficient compared to DeepInsight and is scalable to large datasets. In terms of run time, DeepInsight required 504.8±148.8 seconds (averaged across all datasets) to transform scRNA-seq data into images, whereas Fotomics could complete the imagification process in 63.7±43.7 seconds, expediting the transformation by over 7.9 times in average (dataset-specific runtime and hardware specification is detailed in **Figure 6**). DeepInsight relies on t-SNE dimensionality reduction, which is intensive computationally compared to the highly efficient FFT transformation in Fotomics, making our proposed method easy to scale.

**Figure 6.**
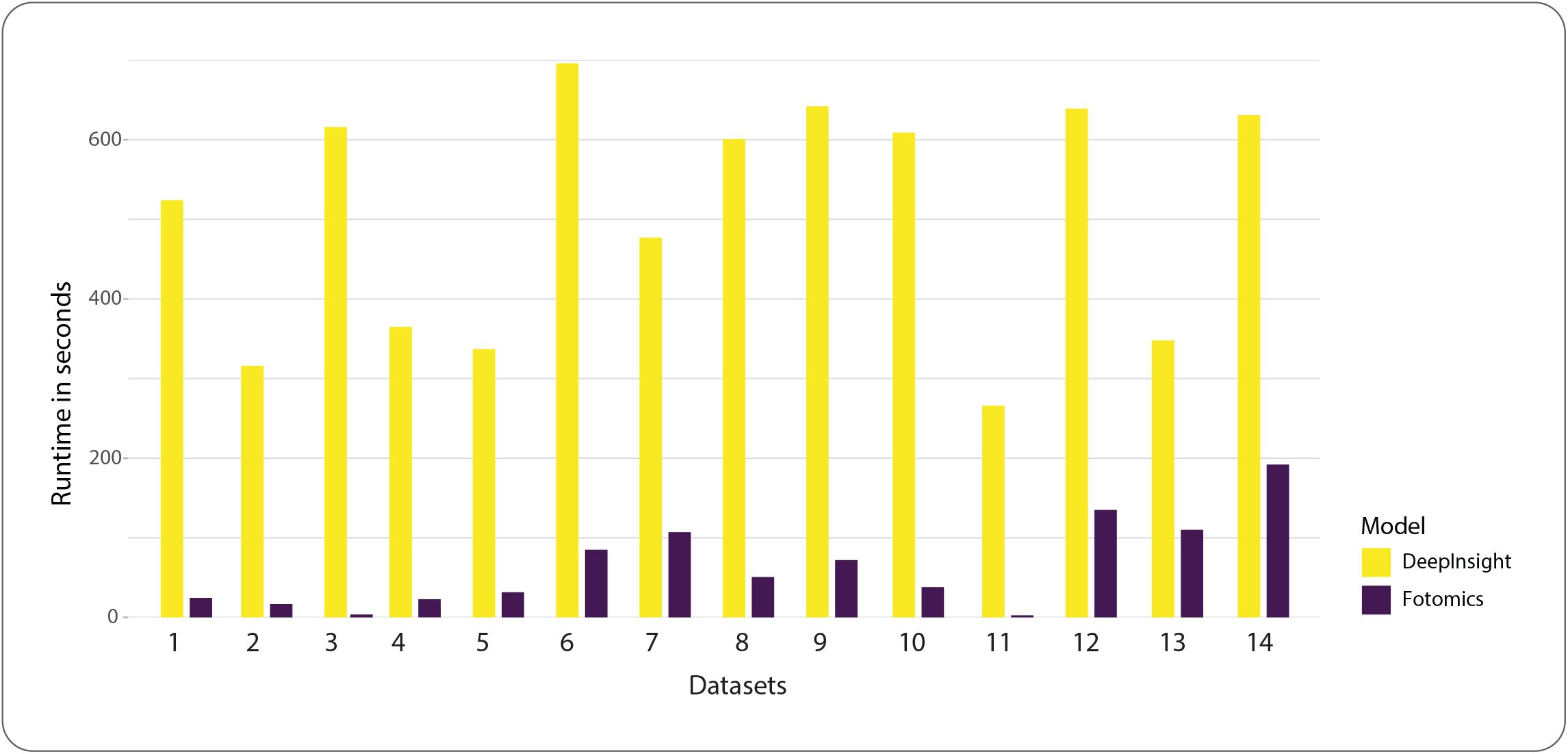
Run time in seconds for converting scRNA-seq profiles of all cells into the corresponding images using Fotomics and DeepInsight imagification techniques. The run time is reported for each dataset. Dataset numbers correspond to the numbers in Table 1. Hardware specifications include Platform: Google Colab; GPU: 1xTesla K80, 2496 CUDA cores, 12GB GDDR5 VRAM; CPU: single-core hyper-threaded Xeon Processors (one core, two threads).

In terms of the memory usage, both methods are comparable (Fotomics: 2.3k MiB and DeepInsight: 2.2k MiB, averaged across all datasets). Memory usage is dependent on the size of the dataset as both methods load the dataset into memory. In fact, it is essential for DeepInsight to load the entire dataset into memory to estimate t-SNE embeddings. However, Fotomics can, in principle, convert samples to images one at a time (with the cost of increased execution time) and can be optimised accordingly when insufficient memory is an issue.

Single-cell omics datasets often represent a heterogenous and imbalanced cell population within a tissue wherein the majority of cells come from a few classes. The average accuracy or F1-score is often biased towards the performance of the majority class and masks performance variations in predicting rare cell types or smaller cell subpopulations. It is, therefore, essential to investigate the performance of models in predicting each cell type rather than merely reporting the aggregated performance on the entire test set. Hence, to assess the performance of Fotomics on each cell type within the test set (20% holdout samples), we reported in **Figure 7** the confusion matrices of selected datasets, i.e., ‘deng read’ (number of cells = 268, number of cell types = 6), ‘pancreas FACS (1961, 10), ‘yan’ (90, 6), and ‘zeisel’ (3005, 9). The results clearly support the utility of Fotomics in classifying small subpopulations of cells. Interestingly, although misclassifications were observed in some cell types, the predicted labels and the actual cell types are often within the same cellular category (e.g., misclassification of *pancreatic A* and *pancreatic D* cells in ‘pancreas FACS’ data set), further demonstrating the power the proposed method.

**Figure 7.**
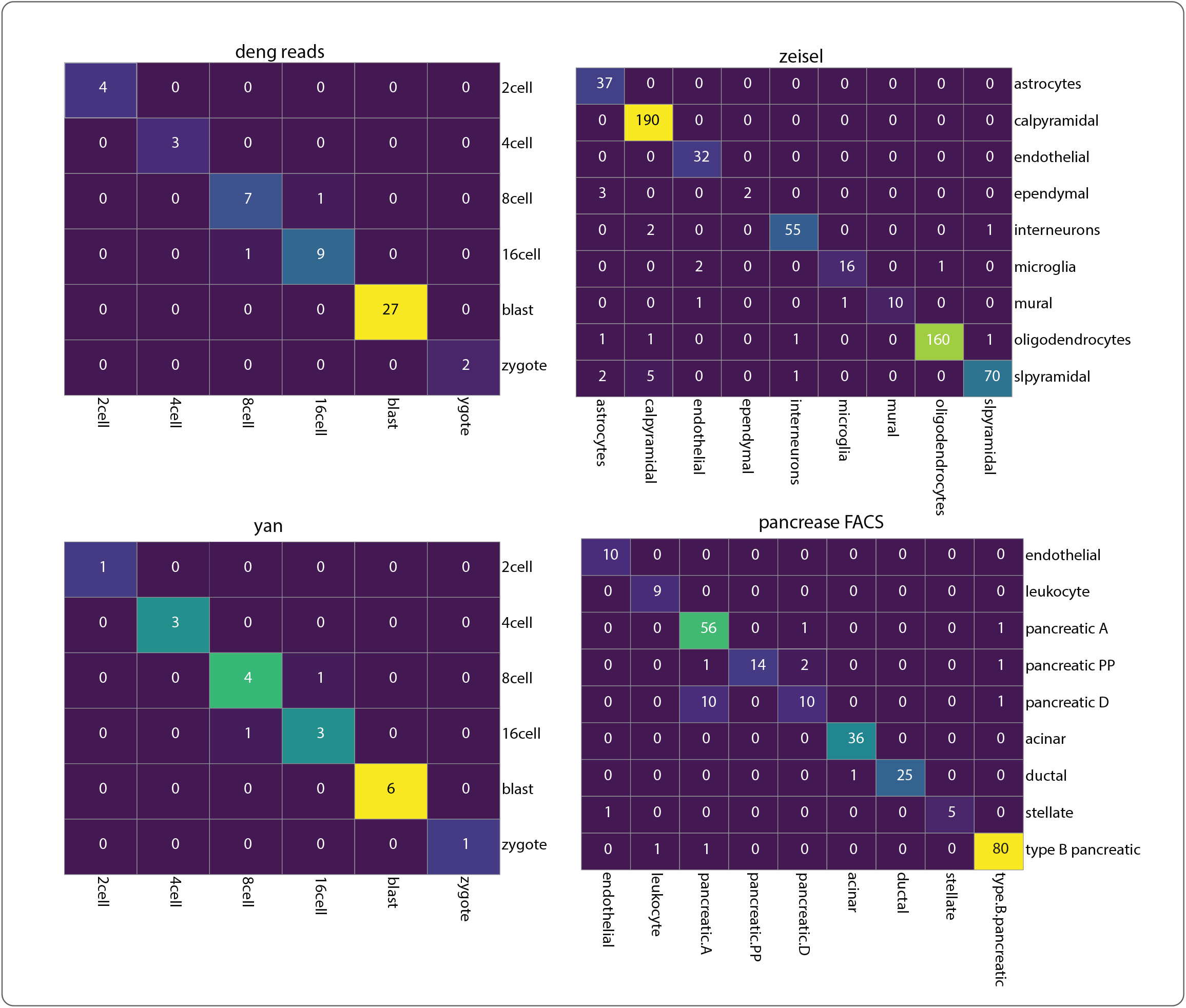
Confusion matrices detailing the performance of CNN classification on the test set (20% randomly held out cells) on selected scRNA-seq datasets (see Table 1 for dataset details). Rows represent annotations (i.e., true classes), while columns represent predictions. Confusion matrices report the proportion of false positives, false negatives, true positives, and true negatives, allowing a more detailed analysis of cell-specific miss-classification.

We further applied the Fotomics imagination to single-cell epigenomics profiles obtained via scATAC-seq (Assay for Transposase-Accessible Chromatin using sequencing). We used the Leukemia scATAC-seq dataset comprising a mixture of 6 cell types, including monocytes and lymphoid-primed multipotent progenitors isolated from a healthy human donor, and leukemia stem cells (x2) and blast cells (x2) isolated from two patients with acute myeloid leukemia [31]. Data were preprocessed as previously explained [32] [ref]. Fotomics imagification combined with CNN significantly outperformed traditional classifiers (accuracies on 25% holdout test set are: Fotomics: 72%, SVM: 46%, RF: 48%, KNN:52%). While Deepsight imagification resulted in higher accuracy (accuracy: 0.845), Fotomics was an order of magnitude faster than DeepInsight in converting scATAC-seq profiles (a total of 352 cells) into the corresponding images (Fotomics: 8.02 seconds, DeepInsight: 4 minutes and 9 seconds). Together, our results demonstrate the utility of the proposed method in converting single-cell omics to images for CNN-based cell identification.

## Conclusions and Future Directions

Depending on the underlying technology used, various omics profiles can include measurements of several hundred to thousands of molecules. The term *omics imagification* is proposed in this paper as a process of converting a vector representing these numerical measurements into an image with a one-to-one relationship to the corresponding sample. The proposed imagification approach transfers omics features such as gene-expression vectors to two-dimensional images to classify the sample phenotype using automated image-recognition technology. The proposed method, called *Fotomics* (*<u>F</u>ourier transformation-based <u>omics</u>*) imagification, relies on the fast Fourier transformation process to efficiently map associated genomics features into a two-dimensional Cartesian plane to generate the corresponding images. Such an image conveys more information about cell types into the classifier, including gene expressions as well as gene-gene relationships obtained from the entire omics profile. The latter enable systems-level and holistic representation of omics profile into an image for pattern recognition using CNN models. The proposed method was applied to multiple single-cell RNA sequencing (scRNA-seq) datasets for evaluation. The experimental results on single-cell RNA-sequencing profiles demonstrate that a simple CNN architecture can significantly outperform commonly used machine learning techniques such as RF, KNN, and SVM. Fotomics was performed competitively well compared to a former state-of-the-art image conversion method, DeepInsight, in terms of prediction power and significantly outperformed DeepInsight in terms of computational runtime.

In this study, the utility of Fotomics was evaluated for automated cell identity mapping via supervised classification of omics images using a convolutional neural network. In principle, omics images can be further used to conduct other essential analyses of single-cell omics data, such as novel cell type identification [33], developmental trajectory inference [34], and unsupervised clustering [35]. For instance, CNN-based autoencoders can be adapted to extract cell embedding (i.e., a low-dimensional vector of latent features) from single-cell omics images. As demonstrated previously [33], the cell embedding can be used to map novel cells to the closest cell type on a share embedding space generated using a heterogeneous meta-dataset where cells from similar cell types are embedded close to each other and cells from different cell types are embedded far away. Hence, the imagified omics profiles have the potential to enable the discovery of novel cell types. Nonetheless, the utility of extracting cell embedding from scRNA-seq images rather than conventional scRNA-seq vectors for applications beyond cell classification requires experimentations and benchmarking analyses which can be considered as a future direction of this study.

Furthermore, while our proof-of-principle experiment corroborates the efficiency and applicability of Fotomics in converting scATAC-sed data to images for CNN-based cell-type classification, further investigations are required to rigorously support the utility of Fotomics on single-cell omics profiles beyond transcriptomics data.

## Data and Code Availability

All the datasets used in this study are public and accessible from Gene Expression Omnibus (GEO). The entire code base, including the python implementation of the proposed method and compared techniques, are available at https://github.com/VafaeeLab/Fotomics-Imagification

